# A Systems-Level ODE Model of B-Oxidation: Algorithmic Flux Optimization and Simulation of MCADD Pathophysiology

**DOI:** 10.1101/2025.08.15.670634

**Authors:** Ethan Shaw, Henry Chuo

**Affiliations:** Children’s Nutrition Research Center, Department of Pediatrics, Baylor College of Medicine, Houston, TX, USA; Department of Biomedical Engineering, University of Houston, Houston, TX, USA

## Abstract

We introduce a computational model that simulates β-oxidation under varying metabolic conditions by solving a system of ordinary differential equations (ODEs) that describe the flux of key intermediates and cofactors. The model captures the stepwise breakdown of saturated, even-chain fatty acids, incorporating dynamic cofactor coupling and product drainage via infinite sinks to maintain numerical stability and biological realism. An optimization routine based on Nelder-Mead simplex refines initial conditions to minimize system residuals. When simulating medium-chain acyl-CoA dehydrogenase deficiency (MCADD), the model predicts metabolite accumulation upstream of the enzymatic block and downstream depletion – highlighting the sensitivity of flux distributions to enzymatic constraints. This work frames kinetic modeling as a tractable method to interrogate systems-level metabolic imbalance and offers a modular framework for future expansions.

## Introduction

β-oxidation is a mitochondrial pathway that supports ATP synthesis via sequential cleavage of acyl-CoA substrates, particularly under fasting or exercise-induced states. The process begins with acyl-CoA activation in the cytoplasm, consuming ATP and producing AMP, before transport into the mitochondrial matrix through the carnitine shuttle. Inside the mitochondrion, β-oxidation proceeds through a cyclic sequence that generates acetyl-CoA, NADH, and FADH_2_, which feed into the electron transport chain for further ATP synthesis [1].

The biological significance of FAO lies in its role as a flexible energy source, especially under glucose-limiting conditions. Dysregulation of FAO – due to genetic mutations, enzyme deficiencies, or mitochondrial dysfunction – can lead to severe metabolic crises such as MCADD [4]. Quantitative modeling offers a means of simulating these perturbations by capturing transient imbalances and compensatory shifts in cofactor usage [2,5].

The goal of this study is to demonstrate how β-oxidation’s fixed enzymatic steps and constrained stoichiometry can be modeled dynamically. Using ODE-based simulations, we interrogate both normal and pathological states, focusing on MCADD as a case study. While deliberately simplified to emphasize core pathway mechanics, this model establishes a foundation for modular extensions that can incorporate regulatory features in future work [6,7].

## Methods

### Model Construction

The β-oxidation pathway was modeled as a simplified kinetic scheme derived from five core enzymatic steps: thiokinase, first oxidation, hydration, second oxidation, and thiolysis. Reaction velocities were defined using mass-action kinetics, linearly proportional to substrate and cofactor concentrations [3]. Cofactors (ATP, FAD, NAD^+^) were treated as variable, enabling dynamic depletion and regeneration. Downstream products were offloaded into infinite sinks (TCA cycle and ETC) for numerical stability [4].

### Reaction Components and Constants

A complete list of enzymes, cofactors, and metabolites is provided in Table 1. Reaction rate constants (k-values) were sourced from the BRENDA enzyme database [6] or inferred from comparable kinetic models [7,10], and are summarized in Table 2. Each enzymatic transformation was associated with an ODE, describing flux propagation across the system (Table 3).

**Table 1.** Components of Beta-Oxidation pathway model from within mitochondrial matrix.

| Components of the β-Oxidation Model | | | |
| --- | --- | --- | --- |
| Type | Code | Name | Function |
| Enzyme | V1, V10 | Thiokinase | Activates fatty acids for β-oxidation (requires ATP) |
| Enzyme | V2, V7, V8 | Acyl CoA dehydrogenase | $A \rightarrow B$ (First oxidation) |
| Enzyme | V3 | Enoyl CoA hydratase | $B \rightarrow C$ (Hydration) |
| Enzyme | V4, V9 | β-Hydroxy acyl CoA dehydrogenase | $C \rightarrow D$ (Second oxidation) |
| Enzyme | V5, V6 | Thiolase | $D \rightarrow E + F$ (Thiolytic cleavage) |
| Cofactor | G | FAD | Electron acceptor (first oxidation) |
| Cofactor | H | FADH<sub>2</sub> | Reduced electron carrier (to ETC) |
| Cofactor | I | NAD<sup>+</sup> | Electron acceptor (second oxidation) |
| Cofactor | J | NADH | Reduced electron carrier (to ETC) |
| Cofactor | K | ATP | Energy donor for fatty acid activation |
| Cofactor | L | AMP | Byproduct of ATP hydrolysis in activation step |
| Substrate | A | Fatty Acyl-CoA | Activated fatty acid |
| Substrate | B | Trans $\Delta^2$ Enoyl CoA | Intermediate (after first oxidation) |
| Substrate | C | β-Hydroxy acyl CoA | Intermediate (after hydration) |
| Substrate | D | β-Keto-acyl CoA | Intermediate (after second oxidation) |
| Substrate | F | Acyl-CoA | Shortened chain for next cycle (n-2 carbons) |
| Product | E | Acetyl-CoA | Product (enters citric acid cycle) |

**Table 2.** Reaction rate constants (K-values) associated with key enzymes in the β-oxidation pathway, defining the kinetics of metabolic transformations.

| Reaction Rate Constants |  |  |
| --- | --- | --- |
| Rate Constant | Value (1/s or 1/mM·s) | Reaction/Enzyme |
| k<sub>1</sub> | 0.05 | Thiokinase (Acyl-CoA synthetase) |
| k<sub>2</sub> | 0.90 | Acyl-CoA dehydrogenase reaction rate |
| k<sub>3</sub> | 1.20 | Enoyl-CoA hydratase reaction rate |
| k<sub>4</sub> | 1.00 | β-Hydroxy acyl-CoA dehydrogenase |
| k<sub>5</sub> | 1.10 | Thiolase (Acyl-CoA Acetyltransferase) |

**Table 3.** Reaction velocities and their corresponding ODEs in the β-oxidation model, describing the dynamic interactions governing metabolite flux.

| Reaction Velocities and ODEs in Beta-Oxidation |  |  |  |
| --- | --- | --- | --- |
| Reaction Velocity | Equation | Corresponding ODE | ODE Equation |
| V1 | $k1 * K$ | $dA/dt$ | $V1 - V2$ |
| V2 | $k2 * A * G$ | $dB/dt$ | $V2 - V3$ |
| V3 | $k3 * B$ | $dC/dt$ | $V3 - V4$ |
| V4 | $k4 * C * I$ | $dD/dt$ | $V4 - V6 - V5$ |
| V5 | $k5 * D$ | $dE/dt$ | $V5 - V7$ |
| V6 | $k5 * D$ | $dF/dt$ | $V6 - V10$ |
| V7 | $k2 * E$ | $dG/dt$ | $-V8$ |
| V8 | $k2 * A * G$ | $dH/dt$ | $V8$ |
| V9 | $k4 * C * I$ | $dI/dt$ | $V9$ |
| V10 | $k1 * K$ | $dJ/dt$ | $-V9$ |
| | | $dK/dt$ | $-V10$ |
| | | $dL/dt$ | $V10$ |

### Initial Conditions

Initial metabolite concentrations were selected from reported mitochondrial metabolomics data [8,9]. All concentrations were expressed in millimolar units to maintain dimensional consistency [7]. These values served as elastic parameters subject to optimization.

### Computational Approach

Numerical integration was performed to simulate metabolite concentrations over time. MATLAB solvers (ODE45) were used with adaptive time-stepping [11]. To refine equilibrium conditions, initial concentrations were optimized using the Nelder-Mead simplex algorithm via FMINSEARCH, minimizing residual system concentration at t = 10 s [14].

## Results

### Time-Course Simulation

Integration of the ODE system yielded directional mass flow through the β-oxidation pathway. While equilibrium was not achieved within the tested time window, metabolite concentrations exhibited expected trends of substrate depletion and product accumulation (Figure 3, Table 5) [10,11].

**Table 4.**
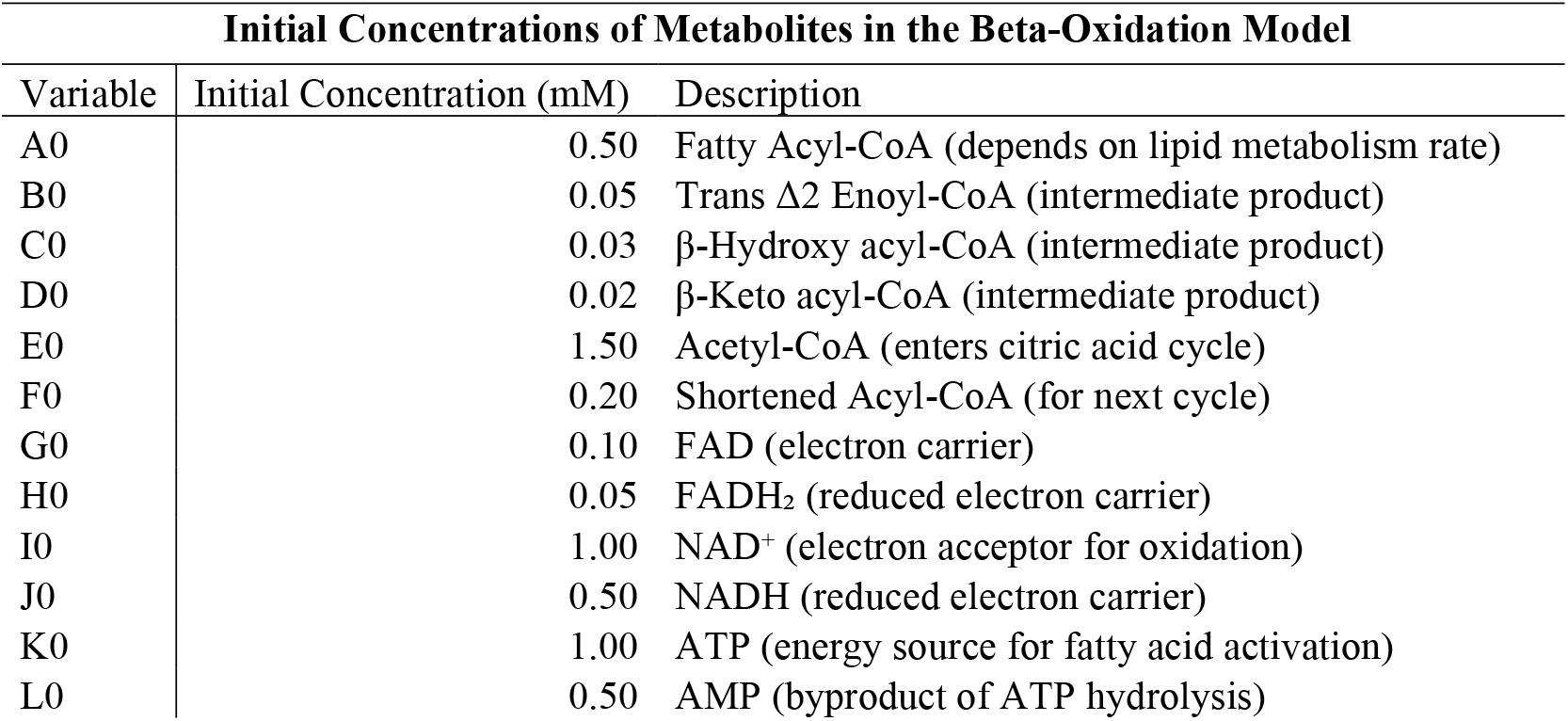
Initial concentrations of metabolites used in the β-oxidation model, defining substrate availability and cofactor levels for testing.

| Initial Concentrations of Metabolites in the Beta-Oxidation Model |  |  |  |
| --- | --- | --- | --- |
| Variable | Initial Concentration (mM) | Description |  |
| A0 | 0.50 | Fatty Acyl-CoA (depends on lipid metabolism rate) |  |
| B0 | 0.05 | Trans $\Delta^2$ Enoyl-CoA (intermediate product) | |
| C0 | 0.03 | β-Hydroxy acyl-CoA (intermediate product) | |
| D0 | 0.02 | β-Keto acyl-CoA (intermediate product) | |
| E0 | 1.50 | Acetyl-CoA (enters citric acid cycle) |  |
| F0 | 0.20 | Shortened Acyl-CoA (for next cycle) |  |
| G0 | 0.10 | FAD (electron carrier) |  |
| H0 | 0.05 | FADH<sub>2</sub> (reduced electron carrier) |  |
| I0 | 1.00 | NAD<sup>+</sup> (electron acceptor for oxidation) |  |
| J0 | 0.50 | NADH (reduced electron carrier) |  |
| K0 | 1.00 | ATP (energy source for fatty acid activation) |  |
| L0 | 0.50 | AMP (byproduct of ATP hydrolysis) |  |

**Table 5.**
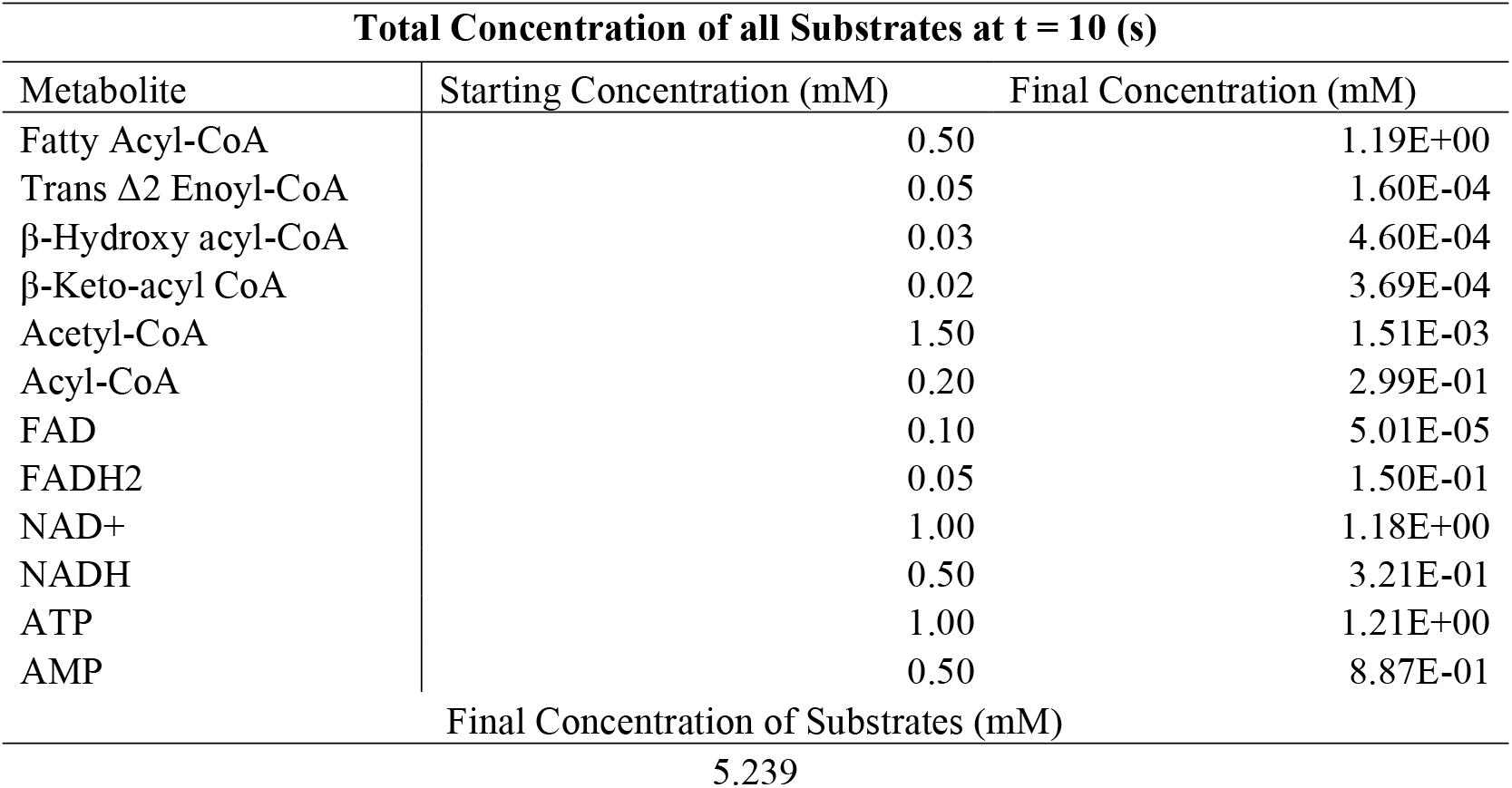
Comparison of initial and final metabolite concentrations, illustrating substrate utilization and system balance at t = 10 (s)

**Figure 1.**
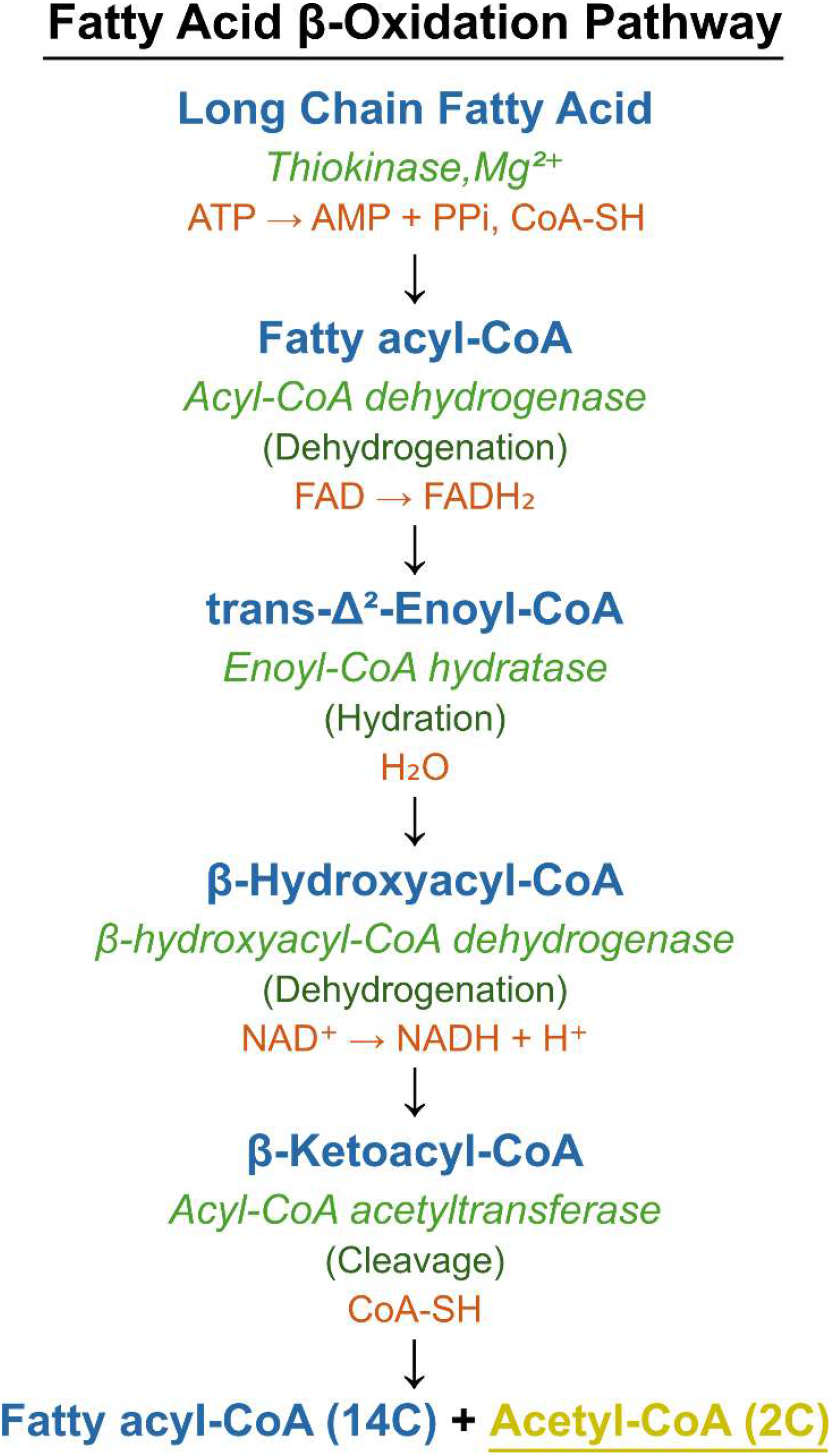
Pathway of Fatty acid Beta-Oxidation from long chain fatty acid to Fatty Acyl-CoA and Acetyl-CoA.

**Figure 2.**
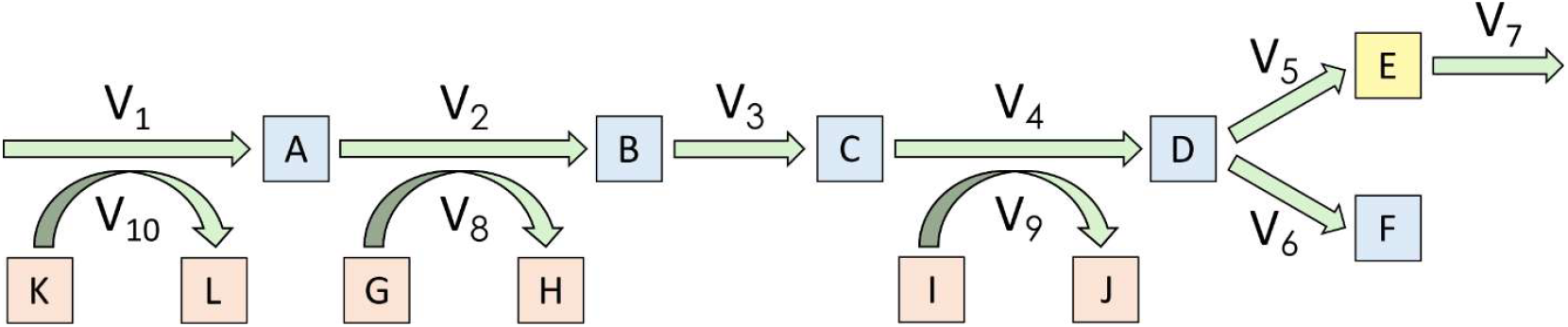
Beta-Oxidation pathway model from within the mitochondrial matrix. Used for equation derivation.

**Figure 3.**
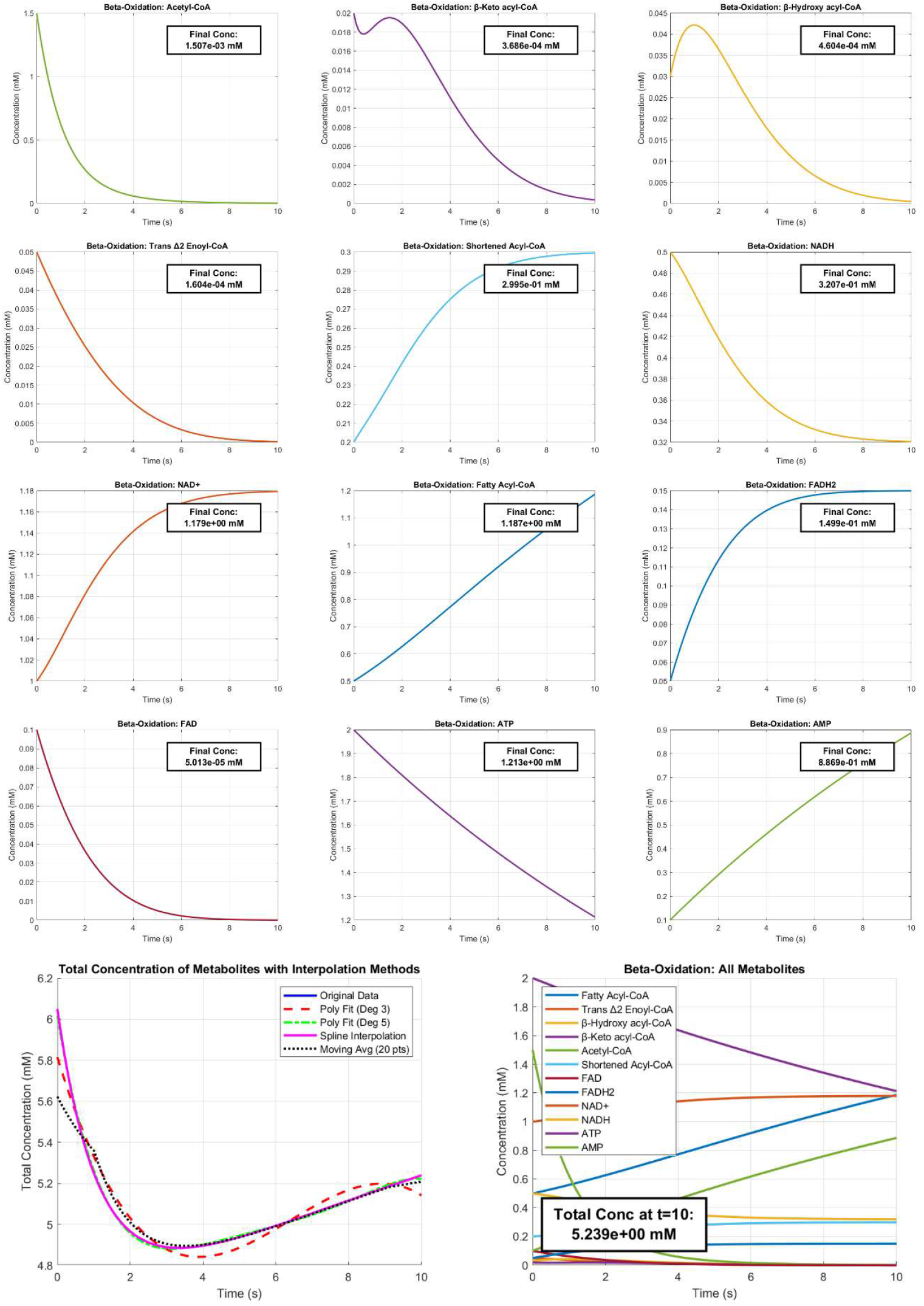
Time-series plots of individual metabolite concentrations and total system concentrations toward t = 10 (s).

**Figure 4.**
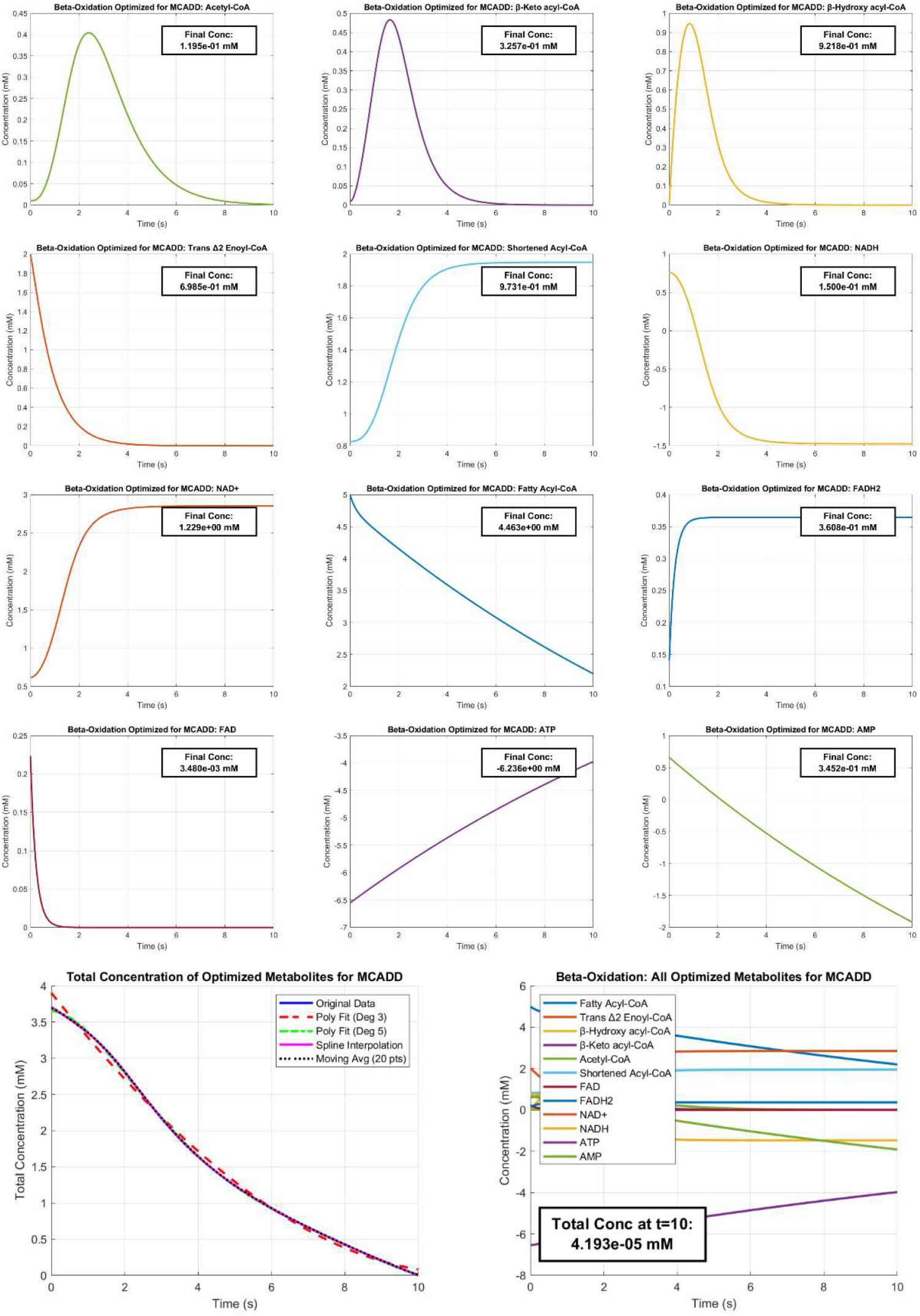
Time-series plots of individual metabolite concentrations and total system concentrations toward t = 10 (s) while optimized for Acyl-CoA Dehydrogenase Deficiency (MCADD) conditions.

### Elastic Refinement

Without optimization, fixed enzyme kinetics and reaction structures prevented complete system balance. Optimization of initial conditions using Nelder-Mead simplex redistributed concentrations, enabling near-zero residual totals at t = 10 s (Table 6) [12,13].

**Table 6.** Comparison of initial, optimized, and final metabolite concentrations, showing how optimization balances fluxes to minimize total concentration at t =10.

| **Comparison of Metabolite Concentrations: Initial, Optimized, and Final State** |  |  |  |
| --- | --- | --- | --- |
| Metabolite | Original Initial Concentration (mM) | Optimized Initial Concentration (mM) | Optimized Final Concentration (mM) |
| Fatty Acyl-CoA | 0.50 | 5.97E-01 | -2.39E-01 |
| Trans $\Delta^2$ Enoyl-CoA | 0.03 | 3.40E-02 | -5.17E-03 |
| β-Hydroxy acyl-CoA | 0.03 | 4.19E-02 | -3.93E-03 |
| β-Keto-acyl CoA | 0.02 | 1.48E-02 | -1.22E-03 |
| Acetyl-CoA | 1.50 | 2.32E+00 | 2.57E-04 |
| Acyl-CoA | 0.20 | 4.86E-01 | 5.71E-01 |
| FAD | 0.10 | 1.09E-01 | 4.00E-02 |
| FADH2 | 0.05 | 7.36E-02 | 1.43E-01 |
| NAD+ | 1.00 | 7.86E-01 | 9.40E-01 |
| NADH | 0.50 | 2.69E-01 | 1.15E-01 |
| ATP | 1.00 | -1.95E+00 | -1.18E+00 |
| AMP | 0.50 | 3.91E-01 | -3.76E-01 |
| Final Concentration (mM) of Substrates (Initial Conditions) at $t = 10$ (s) | | | |
| 5.239 |  |  |  |
| Final Concentration (mM) of Substrates (Optimized Conditions) $t = 10$ (s) | | | |
| -2.25119E-05 |  |  |  |

**Table 7.** Simulation parameters and resulting metabolic effects of Acyl-CoA dehydrogenase deficiency (MCADD), highlighting altered enzyme kinetics, substrate accumulation, intermediate buildup, and disrupted β-oxidation.

| Simulating MCADD: Acyl-CoA Dehydrogenase Deficiency |  |  |  |
| --- | --- | --- | --- |
| Parameter | Current Value | Modified Value (MCADD Simulation) | Effect |
| k<sub>2</sub> (Acyl-CoA Dehydrogenase) | 0.9 | 0.01 – 0.00 | Impaired enzyme function |
| A (Fatty Acyl-CoA) | 0.50 mM | 5.00 mM | Substrate accumulation |
| B (Trans $\Delta^2$ Enoyl-CoA) | 0.05 mM | 2.00 mM | Intermediate buildup |
| C0, D0, E0 (Downstream products) | Normal values | ~0.01 mM | Lack of β-oxidation progression |

### MCADD Simulation

Pathological conditions were modeled by reducing k_2_ (Acyl-CoA dehydrogenase) to near zero. This produced upstream accumulation of fatty acyl-CoA and trans Δ^2^-enoyl-CoA, alongside depletion of downstream intermediates and acetyl-CoA. Flux bottlenecks mirrored clinically observed outcomes of impaired ketogenesis and hypoketotic hypoglycemia [4]. Optimized simulations restored partial system balance by adjusting metabolite pools (Tables 8–9, Figure 5) [2,5].

**Table 8.** Simulation of MCADD disruptions due to reduced k_2_, causing substrate accumulation, intermediate buildup, and impaired β-oxidation. Optimized metabolite concentrations illustrate adjustments restoring balance.

| Optimized Metabolite Concentrations for Combating MCADD |  |  |  |
| --- | --- | --- | --- |
| Metabolite | Original Initial Concentration (mM) | Optimized Initial Concentration (mM) | Optimized Final Concentration (mM) |
| Fatty Acyl-CoA | 5.00 | 5.00E+00 | 2.20E+00 |
| Trans $\Delta^2$ Enoyl-CoA | 2.00 | 2.00E+00 | 1.42E-05 |
| β-Hydroxy acyl-CoA | 0.01 | 1.00E-02 | 1.03E-05 |
| β-Keto-acyl CoA | 0.01 | 1.00E-02 | 2.95E-05 |
| Acetyl-CoA | 0.01 | 1.00E-02 | 1.54E-03 |
| Acyl-CoA | 0.20 | 8.26E-01 | 1.95E+00 |
| FAD | 0.10 | 2.24E-01 | 1.86E-14 |
| FADH<sub>2</sub> | 0.05 | 1.41E-01 | 3.64E-01 |
| NAD<sup>+</sup> | 1.00 | 6.19E-01 | 2.85E+00 |
| NADH | 0.50 | 7.60E-01 | -1.47E+00 |
| ATP | 1.00 | -6.55E+00 | -3.97E+00 |
| AMP | 0.50 | 6.62E-01 | -1.92E+00 |

Final Concentration (mM) of Substrates (Optimized Conditions) at t = 10 (s)
4.19309E-05

## Discussion

This study frames β-oxidation modeling as a tractable computational approach to metabolic disease analysis. By defining enzyme kinetics as inelastic variables and metabolite concentrations as elastic parameters, the model highlights the role of substrate availability in shaping system-level dynamics. Disease states such as MCADD emerge naturally from structural constraints in the ODE framework, producing physiologically consistent bottlenecks [12,13].

While simplifying assumptions (e.g., exclusion of the carnitine shuttle, feedback loops, or isoform-specific kinetics) limit scope, they also provide clarity and computational tractability [7]. The modular structure enables reintroduction of these elements in future iterations [16].

Importantly, the approach demonstrates how computational refinement of initial states can substitute for experimentally inaccessible adjustments in kinetic constants. This emphasizes the diagnostic potential of simulation frameworks for probing disease mechanisms and testing therapeutic strategies in silico [15,17].

## Conclusion

We present an ODE-based kinetic model of β-oxidation that captures both physiological and pathological flux dynamics. Simulations highlight the sensitivity of metabolic pathways to enzymatic impairments, specifically MCADD, and demonstrate how elastic refinement can resolve imbalances. The framework offers a modular, reproducible foundation for future applications in metabolic diagnostics and therapeutic modeling [2,4,12].

## Author Contributions

E.S. and H.C. conceived and developed the computational model. All MATLAB code, figures, and simulations were created independently by the authors. Both contributed equally to the manuscript.

## Data and Code Availability

All source code and a step-by-step protocol are publicly available at: https://dx.doi.org/10.17504/protocols.io.x54v95yepl3e/v2

